# 3-Nitropropionic acid induces histological and behavioral alterations in adult zebrafish: role of antioxidants on behavioral dysfunction

**DOI:** 10.1101/2024.04.29.591507

**Authors:** Melissa Talita Wiprich, Rafaela da Rosa Vasques, Amanda Bungi Zaluski, Kanandra Taisa Bertoncello, Stefani Altenhofen, Darlan Gusso, Gabriel Rodrigues, Adrieli Sachett, Ângelo Piato, Fabio Luiz Dal Moro Maito, Monica Ryff Moreira Vianna, Carla Denise Bonan

**Affiliations:** Programa de Pós-Graduação em Medicina e Ciências da Saúde, Escola de Medicina, Pontifícia Universidade Católica do Rio Grande do Sul, Porto Alegre, RS, Brazil; Programa de Pós-Graduação em Ecologia e Evolução da Biodiversidade, Escola de Ciências da Saúde e da Vida, Pontifícia Universidade Católica do Rio Grande do Sul, Porto Alegre, RS, Brazil; Programa de Pós-Graduação em Biologia Celular e Molecular, Escola de Ciências da Saúde e da Vida, Pontifícia Universidade Católica do Rio Grande do Sul, Porto Alegre, RS, Brazil; Laboratório de Neuroquímica e Psicofarmacologia, Escola de Ciências da Saúde e da Vida, Pontifícia Universidade Católica do Rio Grande do Sul, Porto Alegre, RS, Brazil; Programa de Pós-Graduação em Ciência Biológicas (Bioquímica), Laboratório de Neuroproteção e Doenças Neurometabólicas, Instituto de Ciências Básicas da Saúde, Universidade Federal do Rio Grande do Sul, Porto Alegre, RS, Brazil; Programa de Pós-Graduação em Neurociências, Departamento de Farmacologia, Instituto de Ciências Básicas da Saúde, Universidade Federal do Rio Grande do Sul, Porto Alegre, RS, Brazil; Departamento de Farmacologia, Instituto de Ciências Básicas da Saúde, Universidade Federal do Rio Grande do Sul (UFRGS), Porto Alegre, RS, Brazil; Laboratório de Patologia Bucal, Escola de Ciências da Saúde e da Vida, Pontifícia Universidade Católica do Rio Grande do Sul, Porto Alegre, RS, Brazil; Instituto Nacional de Ciência e Tecnologia em Doenças Cerebrais, Excitotoxicidade e Neuroproteção, Porto Alegre, RS, Brazil

**Author notes:** **Correspondence:** Carla Denise Bonan.

**Keywords:** adult zebrafish, 3-nitropropionic acid, antioxidant, oxidative damage, Huntington’s disease

## Abstract

Huntington’s disease (HD) is a neurodegenerative disease marked by progressive motor and non-motor symptoms such as neuropsychiatric disruption and cognitive dysfunction. It has been reported that some pathogenic mechanisms resulting in neuronal cell death in this disease involve neurodegeneration and oxidative stress. 3-Nitropropionic acid (3-NPA), a natural toxin that promotes the irreversible suppression of mitochondrial complex II, has been used to understand the HD pathogenesis. This neurotoxin mimics the biochemical, central neurodegeneration, peripheral and behavioral phenotype alterations observed in HD. Here we investigated 3-NPA (60 mg/kg) effects on histological and oxidative stress parameters on brain and muscular tissues. We also evaluated the effects of three antioxidant compounds on 3-NPA-induced behavioral phenotypes in adult zebrafish. For the evaluation of the antioxidant effects, adult zebrafish were submitted to a single acute intraperitoneal injection of vitamin C, creatine, or melatonin following 3-NPA chronic administration (60 mg/kg). 3-NPA treatment caused neurodegeneration, but did not alter the muscular tissue. 3-NPA neither change thiobarbituric acid reactive substances (TBARS) nor nonprotein thiol levels. Vitamin C and creatine treatments recovered the hypolocomotion induced by 3-NPA. Also, vitamin C and melatonin treatments improved the memory dysfunction caused by 3-NPA. Altogether, our findings showed that the 3-NPA induces neurodegeneration in adult zebrafish, and the vitamin C, creatine, and melatonin are beneficial in managing HD-like behavioral phenotypes. Thus, these antioxidants could be thought as complementary pharmacotherapies for the treatment of late-stage HD symptoms.

## 1. Introduction

Huntington’s disease (HD) is a complex inheritable neurodegenerative disorder (The Huntington Disease Collaborative Research Group, 1993; Cepeda and Levine, 2022). Clinically, HD is marked by a progressive triad of symptoms, such as motor dysfunction, neuropsychiatric disturbance, and cognitive disruption (Stahl and Feigin, 2020). The motor symptoms are segmented into two stages: in the early stage, it is observed involuntary movements (chorea) mainly in the upper and lower extremities, while the late stage is marked by slow voluntary movements (bradykinesia), stiffness, and motor incoordination (Barry et al., 2022).

This disease is a consequence of an expansion of cytosine-adenine-guanine (≥ 36 repeats) trinucleotide repeats in exon 1 of the huntingtin (*Htt*) gene, which results in a mutation in the *Htt* protein (m*Htt*) (The Huntington Disease Collaborative Research Group, 1993). Similar to *Htt*, m*Htt* is expressed in most human tissues (Bates et al., 2015; Chuang and Demontis, 2021). The m*Htt* causes biochemical alterations including mitochondrial dysfunction, increase in reactive oxygen species (ROS) with subsequent oxidative stress, and consequently neuronal cell death mainly in the striatum (caudate nucleus and putamen) (Gu et al., 1996; Coppen et al., 2018; Sawant et al., 2021). However, biochemical alterations produced by m*Htt* are not restricted to the brain tissue (Chuang and Demontis, 2021).

In this context, the administration of 3-nitropropionic acid (3-NPA) in rodents produces damage in striatal (Mahdy et al., 2014; Milutinovic, 2016; Mehan et al., 2018) and muscular (Hernández-Echeagaray et al., 2011) tissues that are linked to neurobehavioral alterations such as decrease in locomotor activity, gait abnormalities, decrease in strength, and memory deficits (Bortolatto et al., 2017; Wiprich et al., 2020; Sharma et al., 2021; Kumar et al., 2022; Wiprich et al., 2022). These alterations closely mimic the neuropathological and peripherical changes observed in HD patients (Barry et al., 2022).

3-NPA is a powerful inhibitor of the succinate dehydrogenase (SDH; an enzyme located in the mitochondrial complex II of the electron transport respiratory chain), which leads to a reduction in the activity of respiratory chain complex II and a decrease in the energy metabolism (Beal, 1992; Brouillet et al., 1998; Bortolatto et al., 2017). A similar alteration in mitochondrial bioenergetics caused by 3-NPA is detected in *postmortem cerebrum* analysis of HD patients (Morea et al., 2017). The dysfunction in the mitochondrial respiratory chain complex induced by 3-NPA can be the source of reactive oxygen species generation such as thiobarbituric acid reactive substances (TBARS) and reduction in antioxidant defense potentially inducing oxidative stress (Bortolatto et al., 2017; Sindhu et al., 2018; Sharma et al., 2021).

Oxidative stress has been proposed to play a significant role in the pathogenic mechanism of HD (Brondani et al., 2023). In this sense, some studies have investigated the role of antioxidant molecules as a complementary neuroprotective strategy to attenuate the motor and non-motor symptoms of HD (Rebec, 2018; Sindhu et al., 2018; Gunata et al., 2020). Therefore, the 3-NPA pharmacological model represents a valuable tool to elucidate the HD mechanisms and in the search for new pharmacotherapy approaches for this disease.

In the present study, we investigated the 3-NPA effects on histological and oxidative stress parameters on the brain and muscular tissues in adult zebrafish. The protective effect of antioxidants was evaluated on the behavioral alterations induced by 3-NPA in adult zebrafish. We investigated whether 3-NPA administration could induce histological alterations in brain and muscular tissues and brain oxidative stress in adult zebrafish. Additionally, we hypothesized that treatment with antioxidants could recover the behavioral alterations induced by 3-NPA in adult zebrafish.

## 2. Material and Methods

### 2.1. Zebrafish housing conditions

In this study, for all experiments, we used a total of 445 adult zebrafish (*Danio rerio*, AB strain, female/male, 0.3-0.4 g, 6-12 months) from our breeding colony. The fish were maintained in automated recirculating systems (Zebtec, Tecniplast, Italy) with reverse osmosis filtered water equilibrated to reach the essential parameters recommended for the species: temperature (28 °C ± 2 °C), pH (7.0 to 7.5), conductivity (300-700 µS), hardness (80-300 mg/L), ammonia (< 0.02 mg/L), nitrite (< 1 mg/L), nitrate (< 50 mg/L), and chloride levels (0 mg/L). Fish were subjected to a light/dark cycle of 14/10 h, respectively. They were fed with paramecium between 6- and 14-days post-fertilization (dpf), and after 14 dpf, they received commercial flakes (TetraMin Tropical Flake Fish®) three times per day supplemented with brine shrimp (Westerfield, 2000). All fish were manipulated following the Brazilian Council of Animals Experimentation guidelines for Use of Fish in Research (CONCEA) and the Brazilian legislation (11.794/08). The Animal Care Committee of the Pontifical Catholic University of Rio Grande do Sul (CEUA-PUCRS, protocol number 9406/2019) approved the study and registered it in the Sistema Nacional de Gestão do Patrimônio Genético e Conhecimento Tradicional Associado - SISGEN (Protocol No. A3B073D).

### 2.2. Sample size

The sample size for behavioral assays was determined based on the literature according to previous studies performed by our group using zebrafish as an animal model (Wiprich et al. 2023).

The sample size for biochemical assays was established based on the literature according to previous studies using zebrafish as an animal model (Marcon et al., 2019; Valadas et al., 2022).

### 2.3. Chemical compounds

The chemicals 3-nitropropionic acid (3-NPA), creatine, and tricaine were purchased from Sigma (St. Louis, MO, USA). Vitamin C and melatonin were purchased from Bellacqua Manipulation Pharmacy (Porto Alegre, Rio Grande do Sul, Brazil). 3-NPA, creatine, and vitamin C were diluted in saline (0.9% NaCl). Melatonin was diluted in ethanol 0.01%, and tricaine was diluted in the recirculating system water.

### 2.4. Experimental design

Firstly, fish were randomly allocated in a density of three fish per liter into two experimental groups to induction of HD-like phenotypes: saline (vehicle, for the control group) and 3-NPA (60 mg/kg). The animals were maintained and remained in two different aquariums for each experimental group for 29 days.

For behavioral assessments using the antioxidants, the fish from saline and 3-NPA treated groups were selected following randomization and different fish were exposed to the locomotor activity test and the aversive memory task.

Prior to the pharmacological interventions, behavioral (locomotor activity test and aversive memory task), and biochemical analysis the fish did not receive food.

To induction of HD-like phenotypes, fish received chronically seven intraperitoneal (i.p) injections of 3-NPA (60 mg/kg of animal body weight) or saline (vehicle, for the control group) every 96 hours, totaling twenty-four days of treatment. The volume injected was 10-µL. The dose (60 mg/kg) selected was based on a previous study published by our research group, which demonstrated that the 3-NPA chronic administration in adult zebrafish at a dose of 60 mg/kg induces HD-like phenotypes similar to late-stage such as hypolocomotion and memory dysfunction (Wiprich et al., 2020). Both 3-NPA and saline (vehicle, for the control animals) animals remained in the same housing conditions, including water quality, light, and food.

To investigate the effects of antioxidants on the induction of HD-like phenotypes, on day 29 (120 hours after the last injection of 3-NPA and saline) the fish received a single i.p injection of antioxidants, vitamin C (VIT C), creatine (CRE), and melatonin (MEL), or their respective vehicle, 30 minutes before starting each behavioral experiment (locomotor activity test or aversive memory task). VIT C (100 mg/kg), CRE (50 mg/kg), and MEL (10 mg/kg) were administered via i.p in a volume of 10-µL. Doses were selected as described previously by Rodríguez-Martínez et al. (2004) and Marques et al. (2019) using rodents and by Kumari et al. (2019) using zebrafish as animal models. The administration route was based on reported in previous studies performed by our research group using the zebrafish as an animal model (Wiprich et al., 2020; Wiprich et al., 2022). Previously to the induction of HD-like phenotypes and the antioxidants treatment, all fish were weighted and distributed in such a way that the average weight did not differ between the groups.

All chemicals were injected using a 3/10-mL U-100 BD Ultra-Fine™ Short Insulin Syringe with an 8 mm (5/16 in) × 31 G short needle (Becton Dickinson and Company, New Jersey, USA). In all experiments, previously to the injection, fish were anesthetized by immersion in 100 mg/L tricaine solution tamponed with sodium bicarbonate until they showed a lack of motor coordination and a reduced respiration rate. Then, the fish were placed in a sponge soaked in recirculating system water and the injection was performed. Immediately, fish were placed in an aquarium with a highly aerated recirculation system to facilitate their anesthesia recovery. Then, approximately 5 min after tricaine anesthesia, when the fish recovered, they returned to their home aquarium during the period of induction of HD-like phenotypes, or on the day of the antioxidants treatment they were placed in the behavioral test aquarium (Wiprich et al., 2022). The experimental design is shown in Fig.1.

**Fig. 1.**
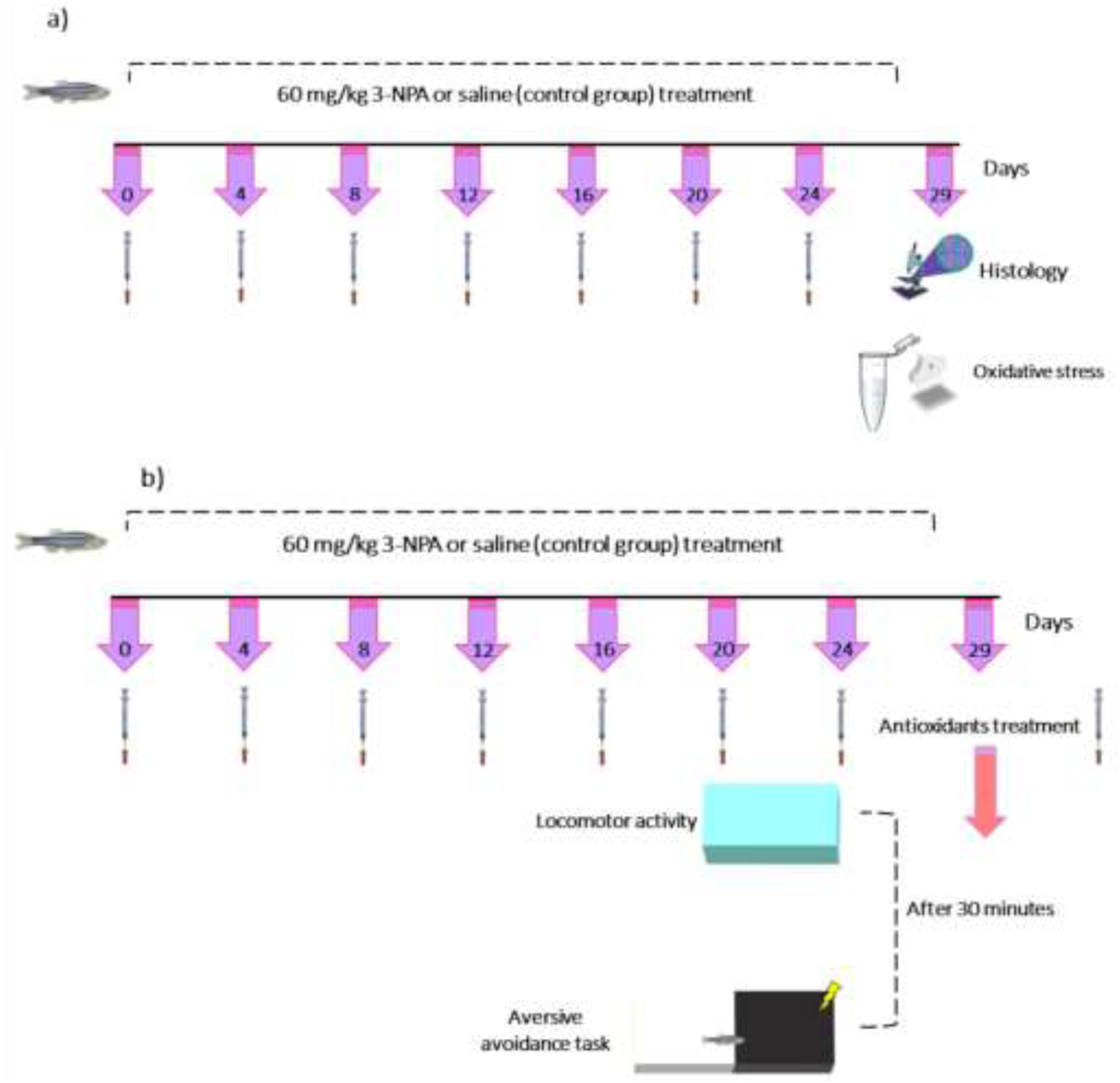
Experimental design of (a) histological analysis, (a) biochemical estimation, and (b) modulation with antioxidants on HD-like phenotypes induced by 3-NPA in adult zebrafish.

### 2.5. Histological analysis

To investigate if the 3-NPA could cause changes in the brain and muscular tissues, we performed the histological analysis in these tissues. Firstly, the fish were cryoeuthanized, and their whole brain and whole muscle were gently dissected. Then, the tissues were put in microtubes (2 mL) containing 10% formaldehyde and remained for 48 h. Subsequently, the muscular tissues were immersed in 17% EDTA for approximately 45 days for bone decalcification. After, all tissues were submitted to a dehydration process through a graded series of alcohol and cleared through a graded series in xylene. Next, the tissues were embedded in paraffin (Patel et al., 2021). The paraffin blocks were sectioned of 4 µm thickness using a rotary microtome (*Lupetec*, MRP-09 model). The tissues were stained with hematoxylin-eosin, examined in an inverted optical microscope (Nikon, SMZ 1500), and analyzed using Image J software. A total of 14 animals were used for the experiments.

### 2.6. Biochemical estimation

To understand the biochemical mechanisms involved in the induction of HD-phenotypes in adult zebrafish and consequently to test one of our hypotheses, we evaluated oxidative stress parameters in adult zebrafish whole brain on day 29 (120 hours) after the last injection of 3-NPA (day 25). Briefly, the fish were euthanized by rapid hypothermic shock (at 2-4 °C), and each whole brain was collected using a scalpel blade. Then, each whole brain was placed in a microtube containing 150 µL of phosphate-buffered saline (PBS, pH 7.4, Sigma Aldrich) per brain. It used a pool of six whole brains per microtube, totaling n=184 animals. The microtube remained on ice until all samples were collected. After, the samples were gently homogenized on ice and centrifuged at 3,000 *g* for 10 min at 4 °C. The supernatants were transferred to microtubes and kept at - 80 °C until the analysis (Marcon et al., 2019; Valadas et al., 2022).

The oxidative stress parameters evaluated were thiobarbituric acid reactive substance (TBARS) and non-protein thiol (NPSH). The analysis of all parameters was based on a previous study carried out by Marcon et al. (2019) and previous protocols detailed by Sachett et al. (2020).

#### 2.6.1. TBARS determination

The lipidic peroxidation was analyzed by the production of TBARS levels. Was added 150 µL of a mixture containing thiobarbituric acid (TBA, 0,5%) and trichloroacetic acid (TCA, 20%, Sigma Aldrich) in the samples (50 µg of protein). Then, samples were heated for 30 min at 100 °C. TBARS levels were determined in triplicate using a microplate reader with an absorbance of 532 nm. The standard used was the malondialdehyde (MDA, 2 mM). The results were expressed as nanomoles (nmol) MDA/mg per protein (Sachett et al., 2020a).

#### 2.6.2. NPSH assay

For analysis, the NPSH levels in the samples (50 µg of protein) were mixture of 50 µL of 6% TCA (Sigma Aldrich). Subsequently, the samples were centrifuged at 10,000 g for 5 min at 4 °C, and the supernatants were collected. In the supernatants was added potassium phosphate buffer (TFK, 1 M) to complete a volume of 245 µL. Next, in all samples was added 5,5’-dithiobis-(2-nitrobenzoic acid) (DTNB, 10 mM). After, the microplate was incubated for 1 h at room temperature in the dark, and then the microplate was read in absorbance at 412 nm. The results were expressed as µmol NPSH/ mg of protein. All samples were performed in triplicate (Marcon et al., 2019; Sachett et al., 2020b).

#### 2.6.3. Protein determination

We determined the protein for all oxidative stress parameters according to the Coomassie blue method using the serum bovine albumin (Sigma Aldrich) as a standard. The samples were read at 595 nm (Marcon et al., 2019).

### 2.7. Behavioral assessments

#### 2.7.1. Locomotor activity test

The locomotor behavior was conducted as previously reported by our research group (Wiprich et al., 2020; Wiprich et al., 2022). The animals were transferred from the maintenance room to the experimental room one day prior to the tests. The experiments were executed between 8:30 and 12:00.. We used 94 fish (n= 8 to 10 per group) and the tests were performed in duplicate. The fish were gently placed individually in experimental glass tanks (30 cm long x 15 cm high x 10 cm wide) containing recirculating system water. The locomotion behavior was recorded for 6 min, and after the 1^st^ min was excluded because it is considered a time to habituation to fish in the novel tank (Wiprich et al., 2022). The records were evaluated using the EthoVision® XT tracking software (version 11.5, Noldus) at a rate of 30 positions per second. The locomotor behavior parameters measured were distance traveled (m) and velocity (m/s, the ratio between distance traveled and movement). The parameter movement was defined as the mobile time when the velocity was higher than 0.6 cm/s. When the animal reached the stop velocity (less than 0.59 cm/s), it was defined as the immobile time (Wiprich et al., 2022).

#### 2.7.2. Aversive memory task

One day prior to the aversive memory task, the fish were moved from the maintenance room to the experimental room. The task was executed (n = 10 to 11 per group) between 8:30 and 12:00 (Blank et al., 2009; Wiprich et al., 2022). Briefly, the apparatus consists of a tank (18 cm long x 9 cm wide x 7 cm high) separated by a mobile guillotine door which divided the tank into two areas of equal size: one black (right side, 8 lux) and one white (left side, 130 lux). The task consists of two sessions denominated training and test sessions, with a 24-h interval between them. In both training and test sessions, each fish was gently placed in the white side of the tank task with approximately 3 cm of recirculating system water and with the mobile guillotine door closed for 1 min to habituation and environmental recognition. After this period, the mobile guillotine door was lifted, and when the fish crossed into the black side of the tank, the mobile guillotine door was closed again, and two electrodes attached to an 8 V stimulator produced when manually activated for 5 s an electric shock (3 ± 0.2 V). The fish was gently removed from the apparatus and placed in its housing tank. After 24 h, the test session was performed, in which the fish were submitted to the same protocol as the training session but without the electric shock. The latency to enter the black compartment during each session was measured in both sessions, and the expected increase in the test session was used as an index of memory retention. A 180-s ceiling was imposed on test session latency measurements.

### 2.8. Statistical analysis

The normality of all data was evaluated by the Shapiro-Wilk test. Locomotor activity test data were analyzed by Two-way ANOVA followed by Bonferroni as a *post-hoc* test was used to analyze the parametric data from the locomotor test, while nonparametric data from the locomotor test was performed and adjusted through the Log-transformation, and thus the data were analyzed using two-way ANOVA followed by Bonferroni as a *post-hoc* test. Aversive memory training and test latencies for each group were compared using the Mann-Whitney *U* matched pairs test.

The data from the aversive memory task were nonparametric and was used the Mann-Whitney *U* test to compare the latencies for each group and to compare the latencies of multiple groups were used the Kruskal-Wallis and Mann-Whitney *U* tests. Also, the Student’s test analyzed oxidative stress data for unpaired samples.

The exclusion criteria were the following: for the locomotor test, the animals were excluded when the software loose more than 10% of the total time spent in the arena, and for the aversive memory task, the animals were excluded when they took longer than 60 s to entry in the dark compartment in the session training.

Locomotor and oxidative stress data are described as mean ± standard error of the mean (S.E.M), while aversive memory task data are described as interquartile range ± median. For all comparisons, a significance level of *p < 0.05* was considered. GraphPad Prism 8 (La Jolla, CA, USA) software was used for statistical analysis.

## 3. Results

### 3.1. Histological analysis

As shown in Fig. 2a, the control group exhibited small regions with neurodegeneration compared with the 3-NPA group, which showed larger regions with neurodegeneration associated with decreased granular cells. However, in both groups, we did not observe inflammation in brain tissue (Fig. 2a). Also, the control and 3-NPA groups did not show histopathological differences in the muscular tissue (Fig. 2b).

**Fig. 2.**
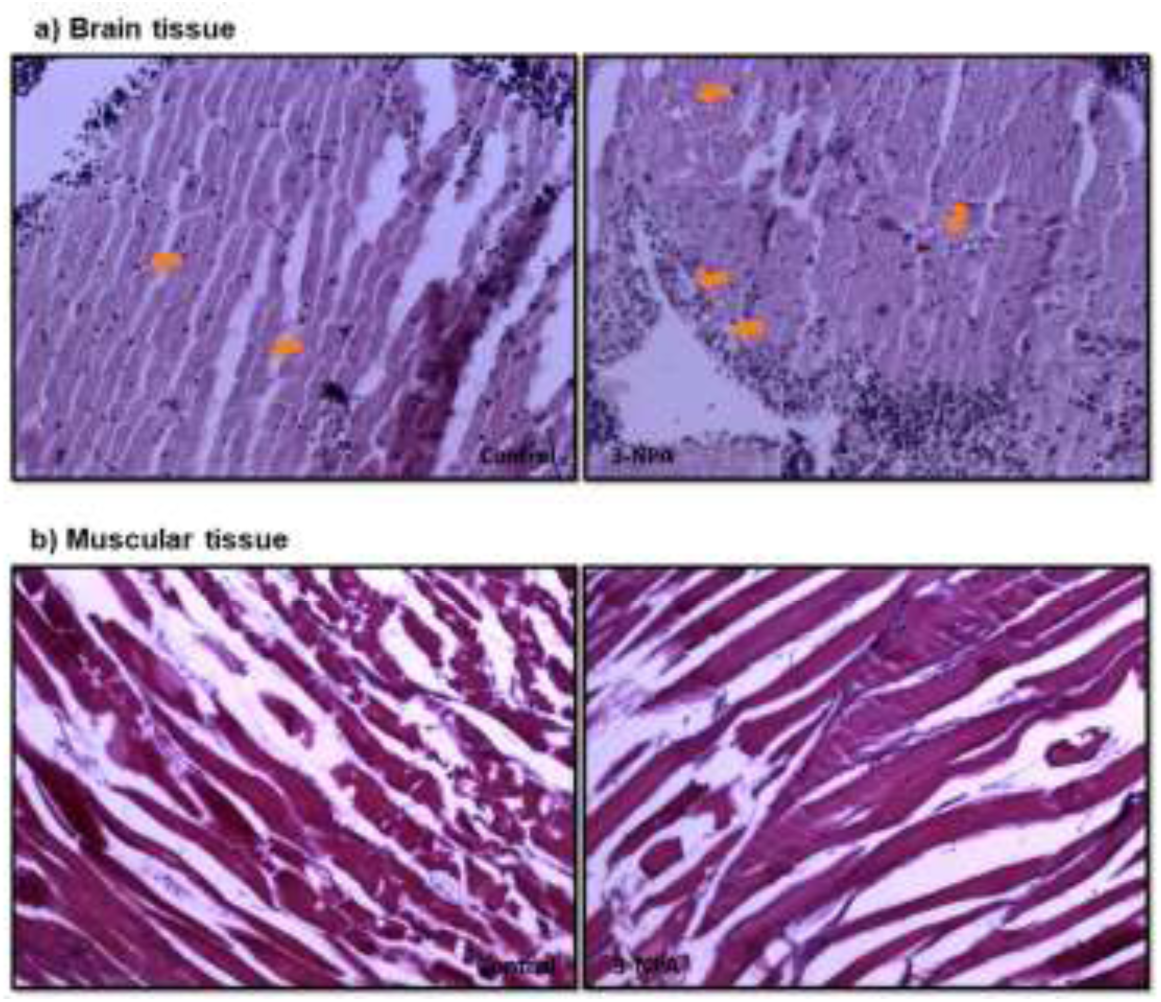
Effect of 3-NPA on (a) brain tissue and (b) muscular tissue. The arrows indicate the regions damage.

### 3.2. Biochemical assays

As depicted in Fig. 3a and Fig. 3b, the 3-NPA treatment did not significantly change TBARS and NPSH levels when compared with the control group (Fig. 3a; *p* = 0.6623, Fig. 3b; *p* = 03583). Also, there were no significant differences between the naïve group (without any i.p. injection) and the control group (TBARS; *p =* 0.5383, NPSH; *p =* 0.8682; data not shown), as well as the naïve and 3-NPA groups (data not shown).

**Fig. 3.**
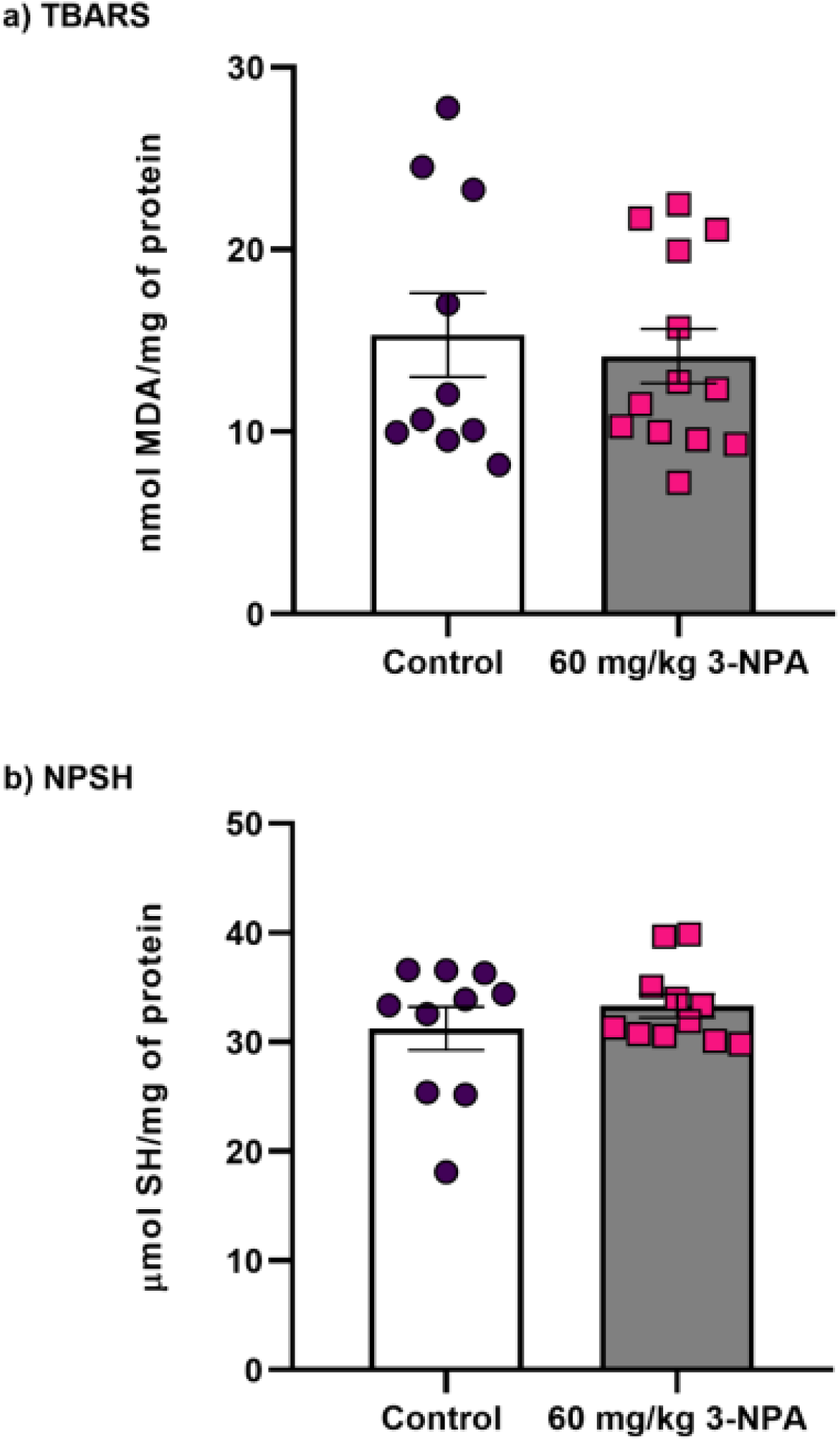
3-NPA effects on (a) thiobarbituric acid reactive substance (TBARS) and (b) non-protein thiol (NPSH) levels. Data are expressed as mean ± S.E.M (*pool* of six whole brains, n=184) and were analyzed by Student’s *t-*test.

### 3.3. Behavioral assessments

The locomotor activity test and aversive memory task were conducted to test whether the antioxidants could recover the hypolocomotion and memory dysfunction HD-like phenotypes induced by 3-NPA.

#### 3.3.1. Locomotor test

As illustrated in Fig. 4a and Fig. 4b, the animals treated with the VIT C and CRE antioxidants recovered the hypolocomotion phenotype (the decrease in distance travelled) induced by 3-NPA (Fig. 4a, 3-NPA, F _(1,35)_ = 14.01; *p =* 0.0007; VIT C, F _(1,35)_ = 2.095; *p =* 0.1567; interaction, F _(1,35)_ = 12.28; *p =* 0.0013; Fig. 4b, 3-NPA, F _(1,36)_ = 5.678; *p =* 0.0226; CRE, F _(1,36)_ = 0.2763; *p =* 0.6024; interaction, F _(1,36)_ = 43.56; *p <* 0.0001). However, the MEL antioxidant treatment did not recover the hypolocomotor alteration induced by 3-NPA (Fig. 4c, 3-NPA, F _(1,50)_ = 13.83; *p =* 0.0005; MEL, F _(2,50)_ = 9.479; *p =* 0.0003; interaction, F _(2,50)_ = 14.11; *p <* 0.0001). Also, CRE (Fig. 4b; *p =* 0.0008) and MEL groups (Fig. 4c; *p =* 0.0002) decreased the distance travelled compared to control animals.

**Fig. 4.**
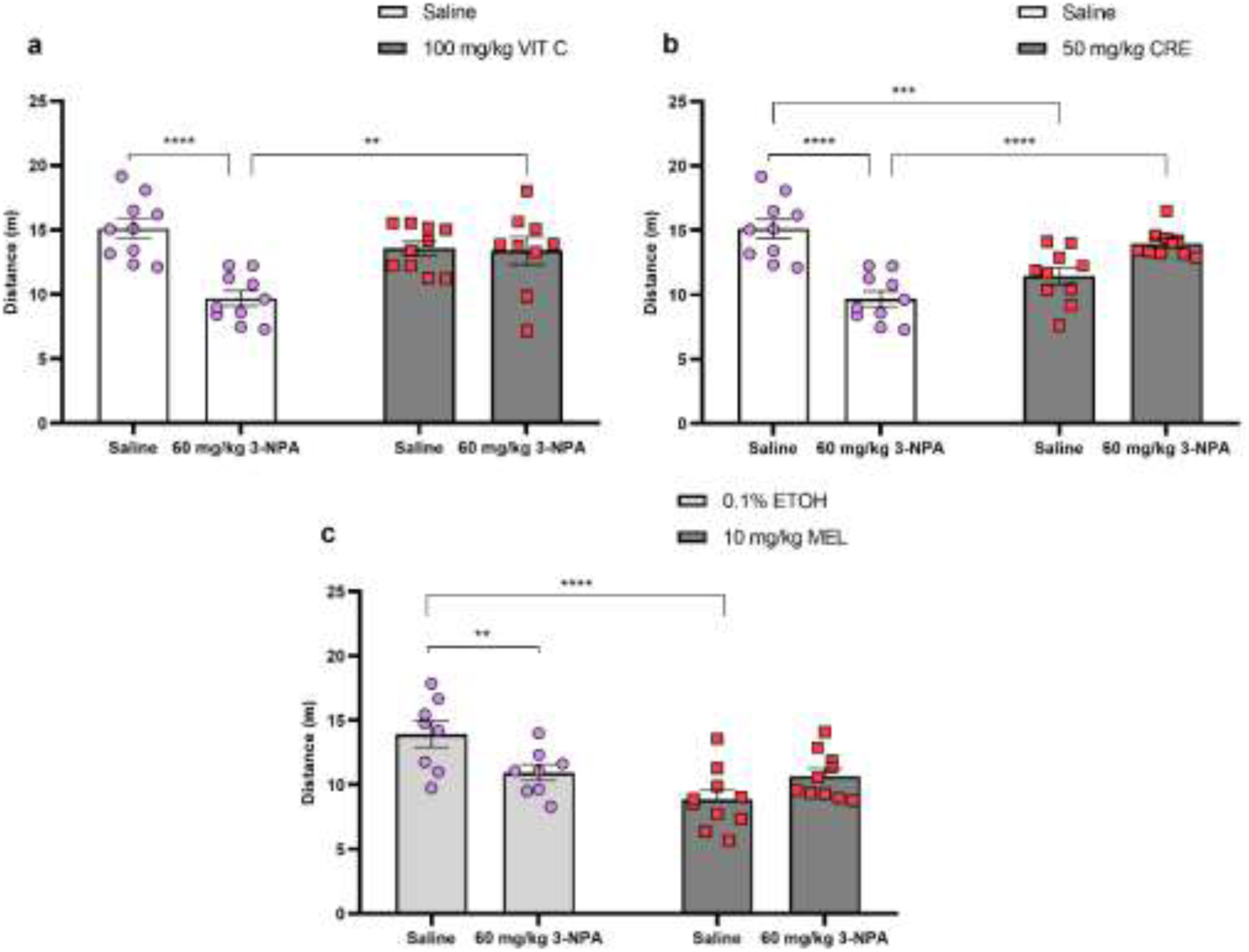
Hypolocomotion phenotype modulation using (a) vitamin C (VIT C; 100 mg/kg), (b) creatine (CRE; 50 mg/kg), and (c) melatonin (MEL; 10 mg/kg) antioxidants. Data are expressed as mean ± S.E.M (n = 8-10 per group) and were analyzed by two-way ANOVA followed by the Bonferroni post-hoc test. ** p < 0.01, *** p < 0.001, **** p < 0.0001.

For all antioxidants, no significant changes were observed in velocity in the 3-NPA group compared to control groups (Supplementary Fig. 1). On the other hand, it was observed that MEL alone caused a reduction in velocity compared to control animals (Supplementary Fig. 1c; *p =* 0.0018).

#### 3.3.2. Aversive memory task

As shown by the difference in latencies between the training and test sessions for each treatment, VIT C (Fig. 5a; *p =* 0.0007) and MEL (Fig. 5c; *p =* 0.0006) recovered the memory dysfunction induced by 3-NPA. In contrast, CRE was not able to recover the 3-NPA-induced memory impairment (Fig. 5b; *p =* 0.2535).

**Fig. 5.**
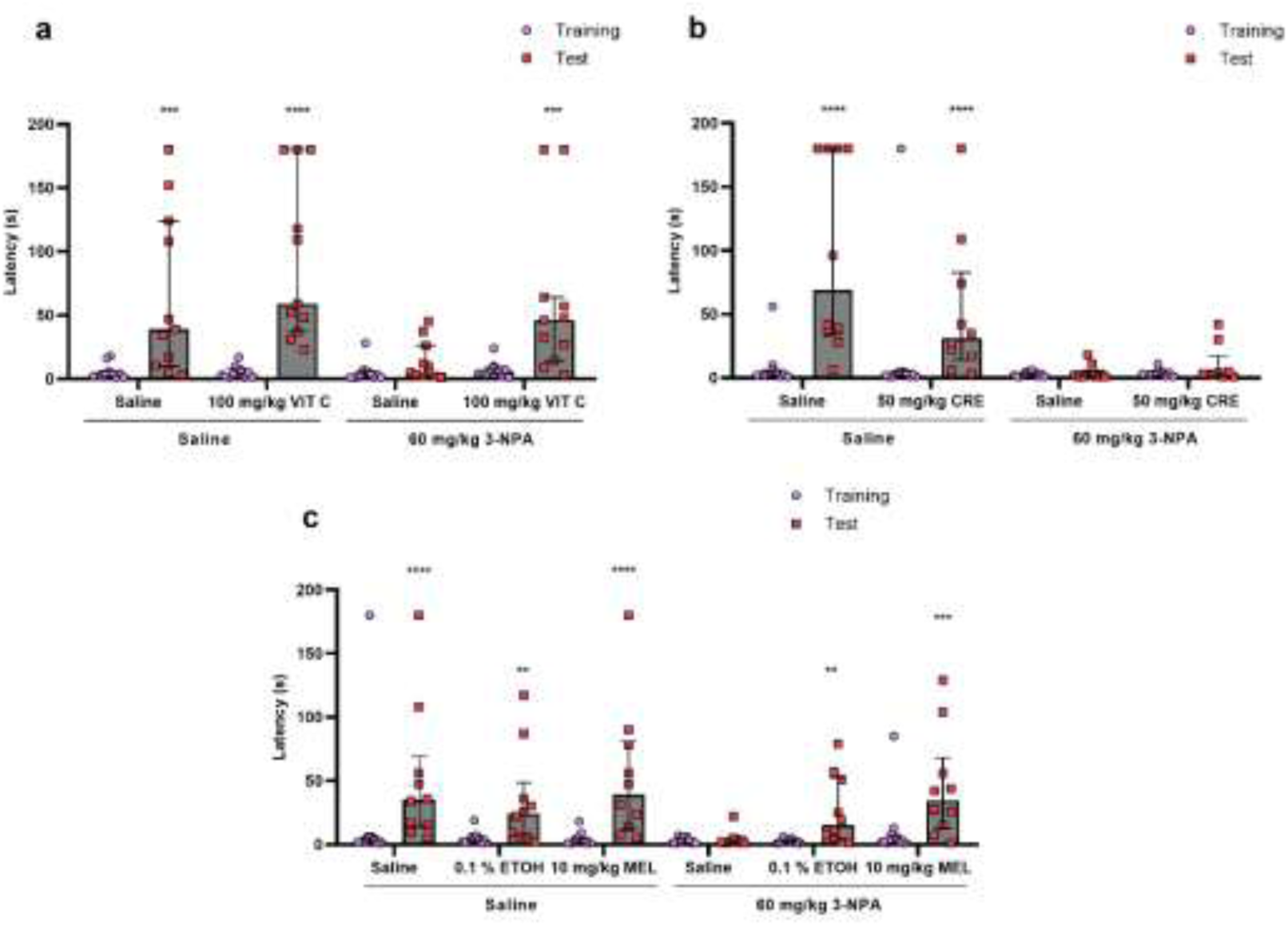
Modulation using (a) vitamin C (VIT C; 100 mg/kg), (b) creatine (CRE; 50 mg/kg), e (c) melatonin (MEL; 10 mg/kg) antioxidants on memory deficits induced by 3-NPA. Data are expressed as median ± interquartile range (n = 10-11 per group) and were analyzed individually per group. ** p < 0.01, *** p < 0.001, e **** p < 0.0001 shows the differences between the training and test sessions for each group analyzed by paired Mann-Whitney *U* test. No significant differences were observed between the training sessions in all groups, as analyzed by the Kruskal-Wallis test.

## 4. Discussion

In this study, it was shown that the 3-NPA induction of HD-like behavioral phenotypes in adult zebrafish did not alter TBARS and NPSH levels. It was also reported that VIT C and CRE recovered the hypolocomotion induced by 3-NPA, and the memory dysfunction was improved by VIT C and MEL.

3-NPA toxin is commonly used to induce HD-like behavioral phenotypes in experimental animal models (Abdelfattah et al., 2020; Wiprich et al., 2020; Moghaddam et al., 2021; Kumar et al., 2022; Wiprich et al., 2022). Behavioral phenotype alterations promoted by 3-NPA include a decrease in locomotor activity, anxiety, and memory dysfunction that are widely associated with the symptoms of advanced (late stage) HD (Bortolatto et al., 2017; Wiprich et al., 2020; Kumar et al., 2022; Salman et al., 2022; Wiprich et al., 2022). In the current study, as expected, 3-NPA induced hypolocomotion and memory dysfunction in adult zebrafish.

It has been previously suggested that motor symptoms of HD are a consequence of central neurodegeneration (Reiner et al., 1988; Kremer et al., 1990; Philips et al., 2015). In this regard, 3-NPA-induced HD animal models usually demonstrate histopathological changes in the brain, including neurodegeneration, neuroinflammation, and gliosis (Mahdy et al., 2014; Dhadde et al., 2016; Mehan et al., 2018; Kumar et al., 2022). Corroborating the literature, we revealed that 3-NPA treatment in adult zebrafish triggered histological brain tissue alteration, such as neurodegeneration and a decrease in the granular cells, but it did not cause neuroinflammation or gliosis. In mammals, the granular cells are mainly located in the hippocampus, which is homologous to lateral pallium in zebrafish (Moser and Moser, 1998; Langova et al., 2020; Gomes-Leal, 2021; Patel et al., 2021).

Further, it has been reported that HD patients and HD transgenic mice show peripheral disturbances, such as muscular skeletal atrophy, resulting from the damage in the muscular tissue caused by *mHTT* (Sathasivam et al., 1999; Ribchester et al., 2004; Mielcarek et al., 2015). We also evaluated the effects of 3-NPA on the histological muscular skeletal tissue and no histopathological changes were observed. In contrast, to the best of our knowledge, the only study evaluating the effects of 3-NPA on the muscular skeletal tissue in rodents described immunohistochemical alterations in muscular skeletal tissue, including muscle atrophy with disorganization in the sarcomere and nuclei of muscle fibers (Hernández-Echeagaray et al., 2011). However, these finding differ from our results. Taken together, our histological analyses in brain and muscular tissue suggest that the motor and non-motor symptoms induced by 60 mg/kg 3-NPA in adult zebrafish may be the result of central neurodegeneration with damage in granular cells.

Previous studies have shown that oxidative stress plays an important role in HD pathogenic mechanisms (D’Edigio et al., 2023). Oxidative stress is caused by an imbalance between the increase in oxidant molecules and their scavenging by antioxidant compounds (Morén et al., 2022). This increase in the oxidant molecules actives the enzymatic (superoxide dismutase, catalase, and glutathione reductase) and non-enzymatic (glutathione, thiols, NPSH, creatine, melatonin, coenzyme Q-10, L-arginine, vitamins C and E, zing, among others) antioxidant strategies to try to counterbalance the oxidants (D’Egidio et al., 2023). The neuronal cells are the most susceptible to oxidative stress because they demand high oxygen and energy consumption and have a lower antioxidant level to combat the oxidant molecules (Morén et al., 2022).

In line with the mentioned above, studies demonstrated an increase in oxidative molecules, such as TBARS, malonaldehyde, and nitric oxide, in the brain of 3-NPA-treated animals (Mehan et al., 2018; Abdelfattah et al., 2020; Habib et al., 2022; Kadir et al., 2022; Brondani et al., 2023). It is also observed an inhibition of antioxidant molecules (superoxide dismutase, catalase, and glutathione) in those animals (Mehan et al., 2018; Kadir et al., 2022; Brondani et al., 2023). Contrary to these findings, we showed that 3-NPA-treated adult zebrafish neither present an increase in the oxidant molecule (TBARS) nor inhibition of non-enzymatic antioxidant NPSH (a subproduct of glutathione) levels in the brain. These divergent findings could be explained because distinct animal species could have different adaptative mechanisms to react to 3-NPA exposure. Therefore, our findings suggest that 3-NPA treatment behavioral and morphological alterations observed in adult zebrafish are more related to changes in neurotransmitter pathways than to an increase in oxidative stress.

Since oxidative stress acts in the HD pathogenesis, studies have investigated the effects of antioxidant compounds as a potential treatment to attenuate disease progression and neurodegeneration (D’Egidio et al., 2023). In this way, we evaluated the effects of three antioxidant compounds (VIT C, CRE, and MEL) on behavioral alterations caused by 3-NPA in adult zebrafish. VIT C and CRE administration significantly recovered the decrease in locomotor activity (hypolocomotion) induced by 3-NPA, while the MEL administration did not recover the hypolocomotion caused by this toxin.

In HD animal models, VIT C administration improved locomotor activity performance compared to control animals (Rebec et al., 2003; Cano et al., 2021). The striatal ascorbate (AA) levels in R6/2 mice (the most used genetic model of HD) showed an increase higher than 50% in WT animals 60 min post anesthesia, while in R6/2 mice showed a decrease ranging from 25% to 50% in AA levels (Rebec et al., 2002). Cortical stimulation led to lower levels of AA release in R6/2 mice compared to WT mice, indicating a dysfunctional link between cortical activation and striatal AA release in HD (Dorner et al., 2009). There is evidence that both R6/2 mice and cells expressing m*HTT* (STHdhQ cells) exhibit an abnormal VIT C flux from astrocytes to neurons and sodium-coupled VIT C transporter deficient (Acunã et al., 2013). Studies have reported that VIT C influences DA synthesis modulating dopaminergic neurotransmission (Hansen et al., 2014; Figueroa-Méndez and Rivas-Arancibia, 2015; Ballaz and Rebec, 2019). In the current study, we found that VIT C administration significantly recovered the decrease in locomotor activity (hypolocomotion) and memory dysfunction induced by 3-NPA. Thus, our findings are in accordance with the literature, suggesting that VIT C has a role in the modulation of locomotor activity and memory possibly by the regulation of neurotransmission systems, such as the dopaminergic system. It should be noted that as far as we know, this is the first study that evaluated the VIT C effects on memory dysfunction induced by 3-NPA.

We also demonstrated that CRE administration recovered the hypolocomotion induced by 3-NPA in adult zebrafish. However, this antioxidant (CRE) did not ameliorate the memory dysfunction induced by this toxin in animals. In two different genetic mice models of HD (R6/2 and N171-82Q), oral CRE supplementation improved the alterations in locomotor activity (Ferrante et al., 2000; Andreassen et al., 2001). Similarly, CRE supplementation improved the motor impairment and the cognitive abnormalities in 3-NPA-treated rodents (Shear et al., 2000; Yang et al., 2009). Also, it was observed neuroprotection in 3-NPA-treated animals supplemented with CRE, where the maintenance of phosphocreatine and ATP levels and an increase in the levels of an oxidative damage marker called 3-nitrotyrosine were observed (Matthews et al., 1998). The CRE supplementation was also tested in some clinical trials. In a randomized multicentric double-blind placebo-controlled study, the CRE administration, at 40 g a day (g/d) for 4 years in early-stage HD patients, worsened chorea symptoms compared to placebo patients. However, the CRE supplementation improved the rigidity component in patients (Hersch et al., 2017). In other two studies, HD patients supplemented with CRE at 10 g/d for 1 year did not show an improvement in their motor and neuropsychological alterations (Tabrizi et al., 2003; Tabrizi et al., 2005). Another clinical trial study observed that CRE treatment did not ameliorate the cognitive dysfunction of HD patients (Rosas et al., 2014). Our data corroborate with previous literature, suggesting that this antioxidant could represent a pharmacotherapy option to attenuate the motor symptoms of late-stage HD. This beneficial effect of CRE on hypolocomotion could be explained by the fact that this antioxidant is metabolized until their phosphorylate form (phosphocreatine). Phosphocreatine could act as an energetic buffer to skeletal muscle, consequently reflecting in the improvement of locomotor activity.

Finally, we observed that MEL administration did not recover the hypolocomotion, while this antioxidant recovered the memory dysfunction induced by 3-NPA in adult zebrafish. Contrary to our findings, most of the results of the previous literature showed that chronic or acute MEL administration reversed the impairment in locomotor activity caused by 3-NPA in HD experimental rat models (Nam et al., 2005; Mu et al., 2011; Tasset et al., 2011; Chakraborty et al., 2014). Another study performed in a *Drosophila* HD model corroborates with the previous studies, indicating that MEL recovered locomotion function (Khyati et al., 2020). MEL treatment in 3-NPA-treated animals also improved memory dysfunction induced by this neurotoxin (Mu et al., 2011). In contrast, other studies have shown that MEL did not reverse the memory dysfunction induced by 3-NPA (Chakraborty et al., 2014). Thus, our findings are in agreement with the described effects for these antioxidants.

Although most of the literature in HD animal models shows beneficial effects of these antioxidants on locomotor activity and memory, it is important to note that there is a scarcity of clinical trials demonstrating the beneficial effects of these antioxidant supplementations in HD patients.

In summary, our data highlight the 3-NPA potential to induce histological changes in the brain, mimicking a histopathological condition (neurodegeneration) as observed in HD patients. Furthermore, while 3-NPA induces histological changes in the brain of adult zebrafish, it does not affect skeletal muscle histology, suggesting that the behavioral alterations observed are likely driven by central rather than peripheral mechanism.The findings also show that VIT C, CRE, and MEL antioxidants could have positive benefits in managing HD-like phenotype symptoms and could be a complementary approach for HD patients in the late-stage of the disease to alleviate motor and cognitive symptoms. However, more clinical trials are needed to confirm the benefits of these antioxidants in HD patients.

## CRediT authorship contribution statement

**MTW:** Conceptualization, performed the experiments, methodology, analyzed and interpreted the data, writing-original draft preparation. **RRV:** Performed the experiments. **ABZ:** Performed the experiments and writing-revised the manuscript. **KB:** Performed the experiments and writing-revised the manuscript. **SA:** Performed the experiments and writing-revised the manuscript. **DG:** Performed the experiments and writing-revised the manuscript. **GR:** Performed the experiments **AS:** Contributed to specific experiments. **AP:** Contributed to conception and design of specific manuscript sections, writing-revised the manuscript. **FLDMM:** Contributed to conception and design of specific manuscript sections, writing-revised the manuscript. **MRMV:** Contributed to conception and design of specific manuscript sections, writing-revised the manuscript. **CDB:** Conceptualization, supervision, funding acquisition, writing-revised, and manuscript editing.

## Ethical statement

All protocols were approved by the Institutional Animal Care Committee from Pontifícia Universidade Católica do Rio Grande do Sul (CEUA-PUCRS, permit number 9406/2019) and Comply with the guideline of the National Council for the Control of Animal Experimentation (CONCEA). This study was registered in the Sistema Nacional de Gestão do Patrimônio Genético e Conhecimento Tradicional Associado – SISGEN (Protocol No A3B073D).

## Acknowledgments

This study was financed in part by Coordenação de Aperfeiçoamento de Pessoal de Nível Superior-Brasil (CAPES)-Finance Code 001, Conselho Nacional de Desenvolvimento Científico e Tecnológico (CNPq; 140290/2019-2; 420695/2018-4; 304450/2019-7), Fundação de Amparo à Pesquisa do Estado do Rio Grande do Sul (FAPERGS; 17/2551-0000977-0), and Instituto Nacional de Ciência e Tecnologia para Doenças Cerebrais, Excitotoxicidade e Neuroproteção.

