## Supplementary Information for "3-Nitropropionic acid induces histological and behavioral alterations in adult zebrafish: role of antioxidants on behavioral dysfunction"

The following figure is related to this article.

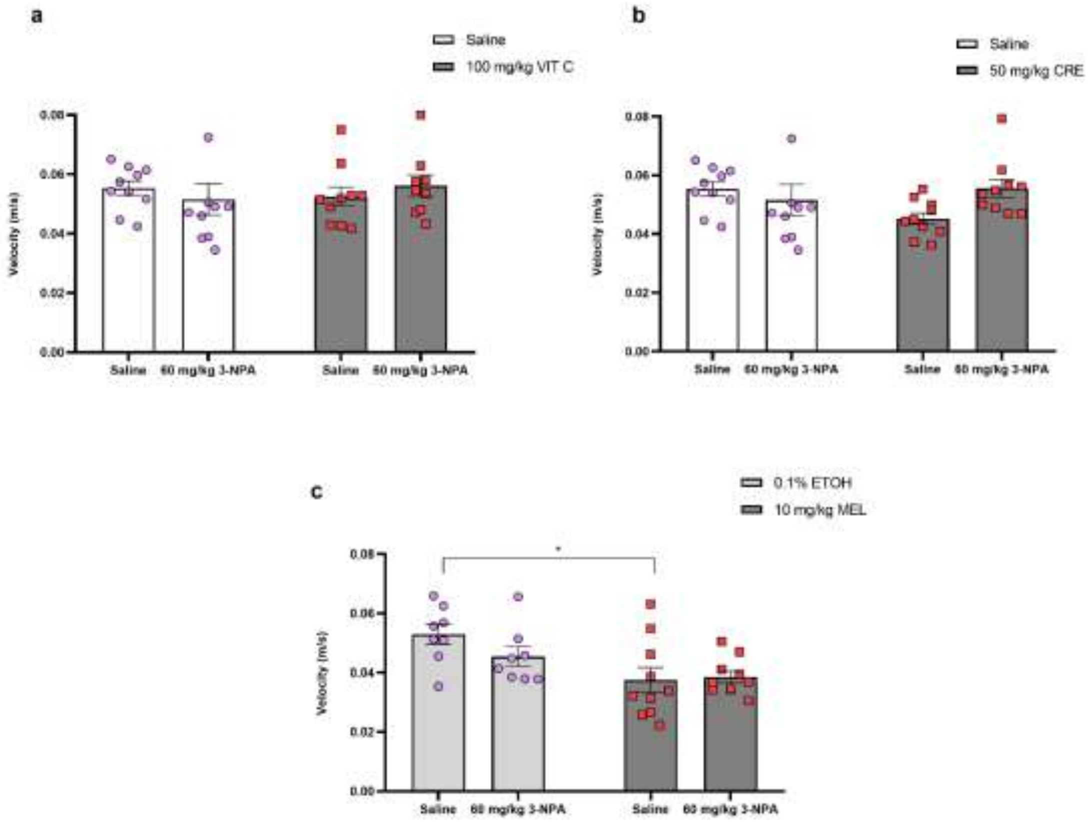


**Supplementary Figure 1.** Effects of antioxidants (a) vitamin C, (b) creatine, and (c) melatonin on velocity after 3-NPA treatment in adult zebrafish. Data are expressed as mean ± S.E.M (n = 12 for each group) and were analyzed by two-way ANOVA followed by a Bonferroni post-hoc test. * $p < 0.05$.
